# Mitochondrial mRNA is a stable and convenient reference for the normalization of RNA-seq data

**DOI:** 10.1101/2021.08.29.458063

**Authors:** Yusheng Liu, Yiwei Zhang, Falong Lu, Jiaqiang Wang

## Abstract

The normalization of high-throughput RNA sequencing (RNA-seq) data is needed to accurately analyze gene expression levels. Traditional normalization methods can either correct the differences in sequencing depth, or correct both the sequencing depth and other unwanted variations introduced during sequencing library preparation through exogenous spike-ins^1-4^. However, the exogenous spike-ins are prone to variation^5,6^. Therefore, a better normalization approach with a more appropriate reference is an ongoing demand. In this study, we demonstrated that mitochondrial mRNA (mRNA encoded by mitochondria genome) can serve as a steady endogenous reference for RNA-seq data analysis, and performs better than exogenous spike-ins. We also found that using mitochondrial mRNA as a reference can reduce batch effects for RNA-seq data. These results provide a simple and practical normalization strategy for RNA-seq data, which will serve as a valuable tool widely applicable to transcriptomic studies.

## Introduction

The normalization of high-throughput RNA sequencing (RNA-seq) data is critical for ensuring the accurate measurement of expression levels since technology-related errors still exist^7-11^. Several normalization approaches have been used to normalize RNA-seq data, referring to the library size concept (such as the approach of Anders and Huber (AH)^12^, trimmed mean of M values (TMM) in the *edgeR* package^10^, and relative log expression (RLE) in the DESeq2 package^13^) or adjusting the distribution of read counts (such as total count (TC), a housekeeping gene count, upper quartile (UQ)^7^, median (Med), quantile (Q)^2^, reads/fragments per kilobase of exon model per million mapped reads (RPKM/FPKM)^14^, and transcripts of exon model per million mapped reads (TPM)^15^). These typical normalization approaches can correct between-sample distributional differences in total read counts^7,10^, and weaken gene-specific effects within samples, including gene length or GC content effects^8,11^. However, they fail to correct for more complex variabilities resulting from library preparation.

Exogenous spike-ins have recently been developed, including the commercial RNA controls raised by the External RNA Controls Consortium (ERCC). ERCC RNA spike-ins consist of a set of polyadenylated transcripts, which mimic natural eukaryotic mRNA and are designed to be added to RNA analysis before or after sample isolation^1-4^. However, the percentage of reads mapped to the ERCC spike-ins could differ from the nominal value and vary widely between technical replicates^6^. In addition, Risso et al.^5^ produced a normalization strategy called remove unwanted variation (RUV), which adjusts for unwanted technical error by performing a factor analysis on suitable sets of control genes (RUVg) or samples (RUVs and RUVr). While the RUV calculation is complicated, RUVg is sensitive to the choice of control genes^5,16^, and RUVs and RUVr cannot work when the treated samples are in different batches from control samples^5^. As such, a better normalization approach with a better reference is needed to analyze RNA-seq data.

Mitochondria are semi-autonomous organelles found in most eukaryotic organisms, and the transcription, processing, and degradation of mitochondrial mRNA (mRNA encoded by mitochondria genome) take place in the mitochondria, independent of nuclear-encoded mRNA^17^. Therefore, we asked whether mitochondrial mRNA could work as an endogenous reference for the normalization of RNA-seq data. In this study, we demonstrate that using mitochondrial mRNA as an endogenous reference is appropriate for normalization of RNA-seq data for inter-sample comparison, which can be a more stable choice than using exogenous spike-ins.

## Results

### Behavior of the mitochondrial mRNA in RNA-seq

To test the feasibility of mitochondrial mRNA as an endogenous reference, we downloaded some published RNA-seq data^18-21^ and calculated the content of mitochondrial mRNA read counts in the total sequencing reads. We found that the mitochondrial mRNA content was similar among different replicates and batches of samples using the same library construction procedures (Fig. 1), although the proportion of mitochondria was different for different samples with the same library construction procedure and the same samples with different library construction procedures (Fig. 1). When we assessed individual genes, we found the mitochondrial mRNA content for each gene was very stable among the different replicates, though it varied among genes (Extended Data Fig. 1). These results suggest that mitochondrial mRNA can be used as an endogenous normalization reference in RNA-seq data.

**Fig. 1.**
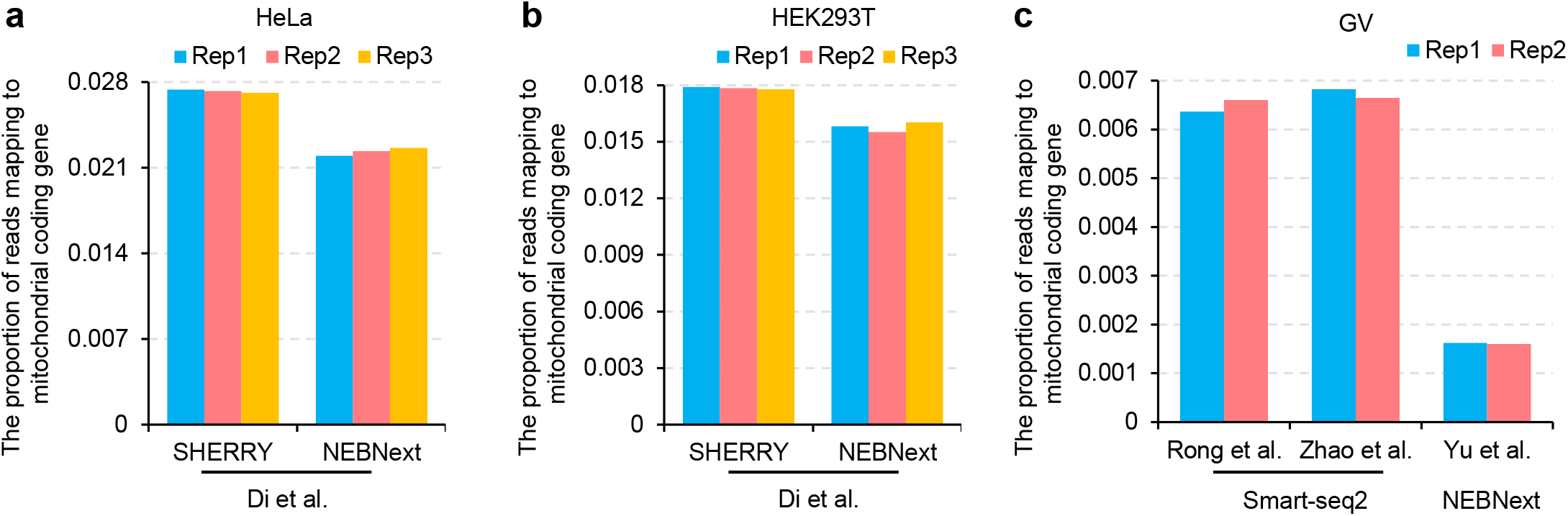
Level of mitochondrial mRNA in multiple RNA-seq datasets. Bar plots indicate proportion of total reads mapped to mitochondrial coding gene in HeLa cells^20^ (**a**), HEK293T cells^20^ (**b**), and mouse GV oocytes^18,19^,21 (**c**). Two or three replicates from the dataset are shown as bar plots. The different methods in constructing the RNA-seq libraries are indicated at the bottom of the plots.

### Mitochondrial mRNA performs more stable than exogenous spike-ins for RNA-seq data normalization

Mitochondrial mRNA is a steady endogenous reference to compare replicates, leading us to assess whether it performs better than exogenous spike-ins for RNA-seq data normalization. Therefore, we downloaded a set of zebrafish control sample data (Con-1, - 2, and -3) from a 2014 article on normalization methods^5^. We found obvious differences in ERCC spike-ins content between the three replicates (Fig. 2a, red vs blue column), which is consistent with the previous report^5^. In comparison, the difference in mitochondrial mRNA content between replicates was slight (Fig. 2b). The big difference in ERCC spike-ins content resulted in large impact on the normalized counts based on the ERCC counts: the linear regression line between any two replicates was far from the diagonal line (y = x, which means the two replicates are identical) (Fig. 2c). Conversely, the linear regression line between any two replicates was close to the diagonal line when we calculated the normalized counts based on mitochondrial mRNA (Fig. 2d).

**Fig. 2.**
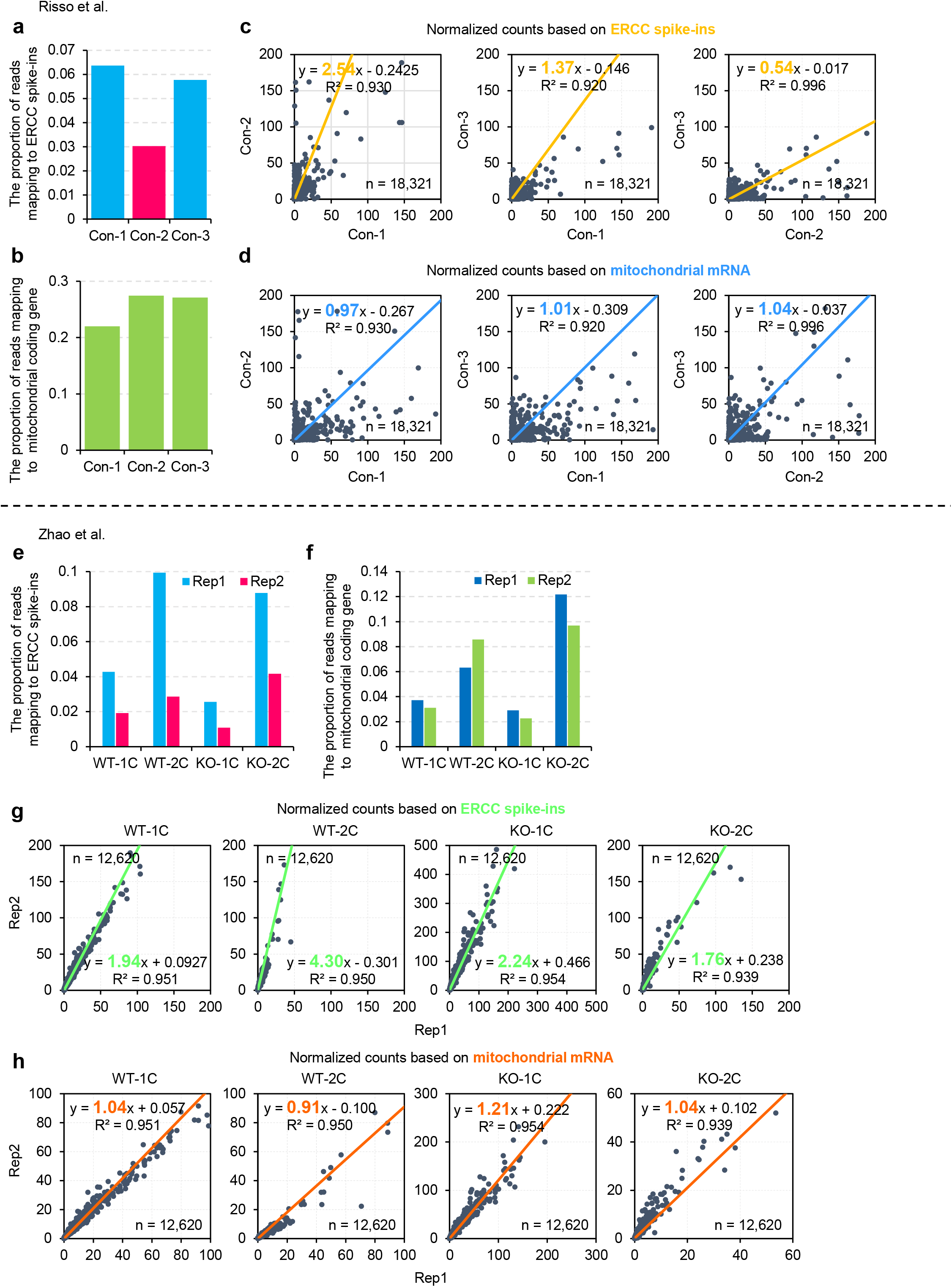
Mitochondrial mRNA performs more stable than exogenous spike-ins in RNA-seq normalization. **a, b**, Bar plots indicate the proportion of total reads mapped to the ERCC spike-ins (**a**) or mitochondrial coding gene (**b**) in a zebrafish control dataset of three replicates^5^. **c, d**, Scatter plots for the counts of genes normalized by ERCC spike-ins (**c**) or mitochondrial mRNA (**d**) between two out of the three zebrafish control replicates. Each dot represents one gene. The number of genes included in the analyses is shown on the bottom right of the graphs. The linear regression equations for the two replicates are shown on the top of each plot with the slope highlighted in yellow (**c**) or blue (**d**). The linear regression line is shown in yellow (**c**) or blue (**d**). **e, f**, Bar plots indicate the proportion of total reads mapped to ERCC spike-ins (**e**) or mitochondrial mRNA (**f**) in a mouse WT and Pabpn1l KO 1C and 2C dataset with two replicates each^19^. **g, h**, Scatter plots for the counts of genes normalized by ERCC spike-ins (**g**) or mitochondrial mRNA (**h**) between the two replicates for each sample of the mouse WT and Pabpn1l KO 1C and 2C dataset. Each dot represents one gene. The number of genes included in the analyses is shown on the top left (**g**) or bottom right (**h**) of the graphs. The linear regression equations for the two replicates are shown on the bottom (**g**) or the top (**h**) of each plot with the slope highlighted in green (**g**) or orange (**h**). The linear regression line is shown in green (**g**) or orange (**h**).

We also downloaded a set of *Pabpn1l* KO data using ERCC spike-ins^19^ and analyzed data from four samples (WT-1C, WT-2C, KO-1C, and KO-2C) with two replicates each. We found that ERCC spike-in content was obviously different between two replicates (Fig. 2e), while the mitochondrial mRNA content was relatively close between any two replicates (Fig. 2f). The linear regression line between any two replicates differed from the diagonal line when ERCC spike-ins were used as a reference for normalization (Fig. 2g), while the linear regression line between any two replicates was close to the diagonal line when mitochondrial mRNA was used as a reference for normalization (Fig. 2h).

These results indicate that mitochondrial mRNA performs more stable than exogenous ERCC spike-ins for normalizing RNA-seq data among replicates.

### Mitochondrial mRNA reduces batch effects in RNA-seq

We analyzed whether using mitochondrial mRNA as a reference can reduce batch effects during RNA-seq analysis by downloading and analyzing different batches of data with ERCC spike-ins from the same laboratory, the same samples (mouse GV oocytes), and the same library preparation procedures^19,21^. We found differences in the content of ERCC spike-ins between different batches (Fig. 3a), reflecting variabilities associated with the addition of exogenous spike-ins to different batches. After normalization with the ERCC spike-ins, slope of the linear regression line between data from two batches was more than 1.5 (Fig. 3b). Conversely, the two batches showed very high parallelism (slope of the linear regression line is close to 1) after normalization with mitochondrial mRNA (Fig. 3c). These results suggested that normalization using mitochondrial mRNA is better than exogenous spike-ins at reducing the batch effect of RNA-seq.

**Fig. 3.**
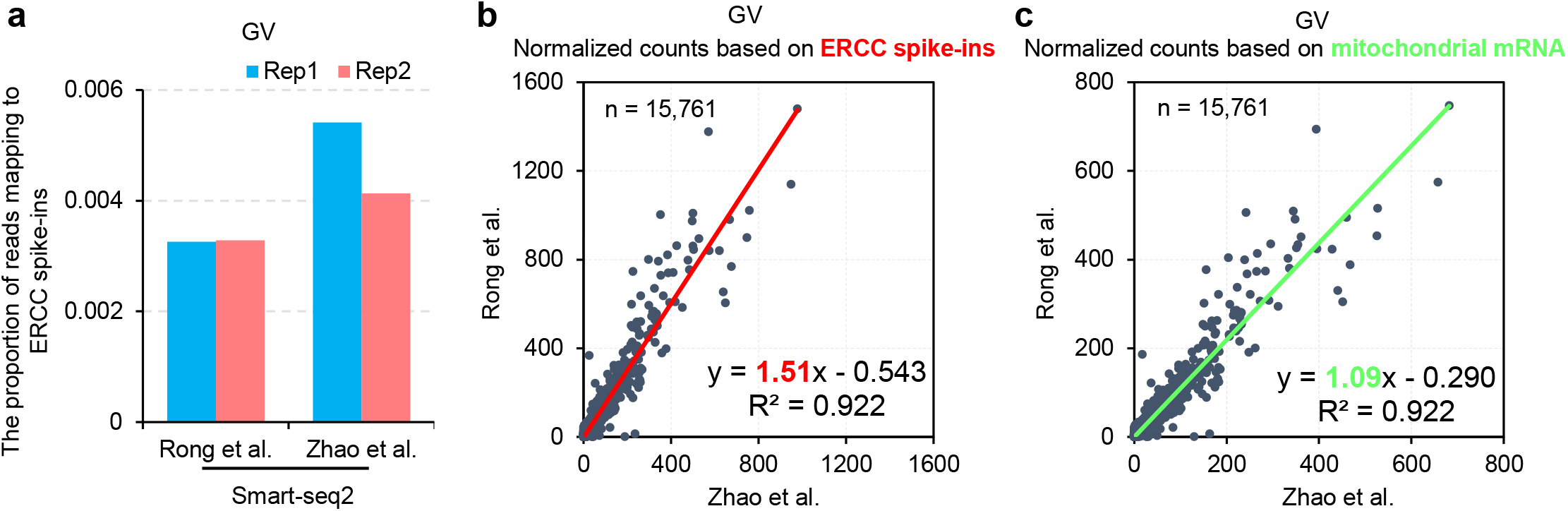
Mitochondrial mRNA can solve batch effects in RNA-seq. **a**, Bar plots indicate the proportion of total reads mapped to the ERCC spike-ins in two mouse WT GV oocyte datasets with two replicates each^19,21^. **b**, Scatter plots for the counts of genes normalized by ERCC spike-ins (**b**) or mitochondrial mRNA (**c**) between the two mouse WT GV oocyte datasets. Each dot represents one gene. The number of genes included in the analyses is shown on the top left of the graphs. The linear regression equations for the two replicates are shown on the top of each plot with the slope highlighted in red (**b**) or green (**c**). The linear regression line is shown in red (**b**) or green (**c**).

### Mitochondrial mRNA as a reference is suitable for differential gene expression analysis

The entire life cycle of mitochondrial mRNA metabolism occurs in the mitochondria, independent of gene regulation in the nucleus^17^. For example, we found that *Btg4* knockout inhibits global poly(A) tail deadenylation of nuclear encoded maternal mRNA, but does not affect mitochondrial mRNA deadenylation in either mouse oocytes or human zygotes (our mouse/human/mito paper). Mitochondrial mRNA is a suitable endogenous reference for RNA-seq data analysis, therefore, we analyzed whether it can be used to analyze differentially expressed genes (DEGs) after treatment that affects gene expression (such as gene knockout). To test this, we analyzed *Btg4* gene knockout data from published datasets^18^ and detected DEGs (p < 0.05, |log2(Fold change)| ≥ 1) normalized by exogenous spike-ins or by mitochondrial mRNA. We found that the thousands of DEGs called through normalization by exogenous spike-ins or by mitochondrial mRNA were highly similar (Fig. 4a) and that their expression fold-change was highly correlated (Fig. 4b, correlation coefficient R^2^ = 1). These results indicate that a normalization strategy using mitochondrial mRNA as a reference is suitable for DEGs analysis for gene knockout samples as long as gene manipulation doesn’t affect mitochondrial mRNA metabolism.

**Fig. 4.**
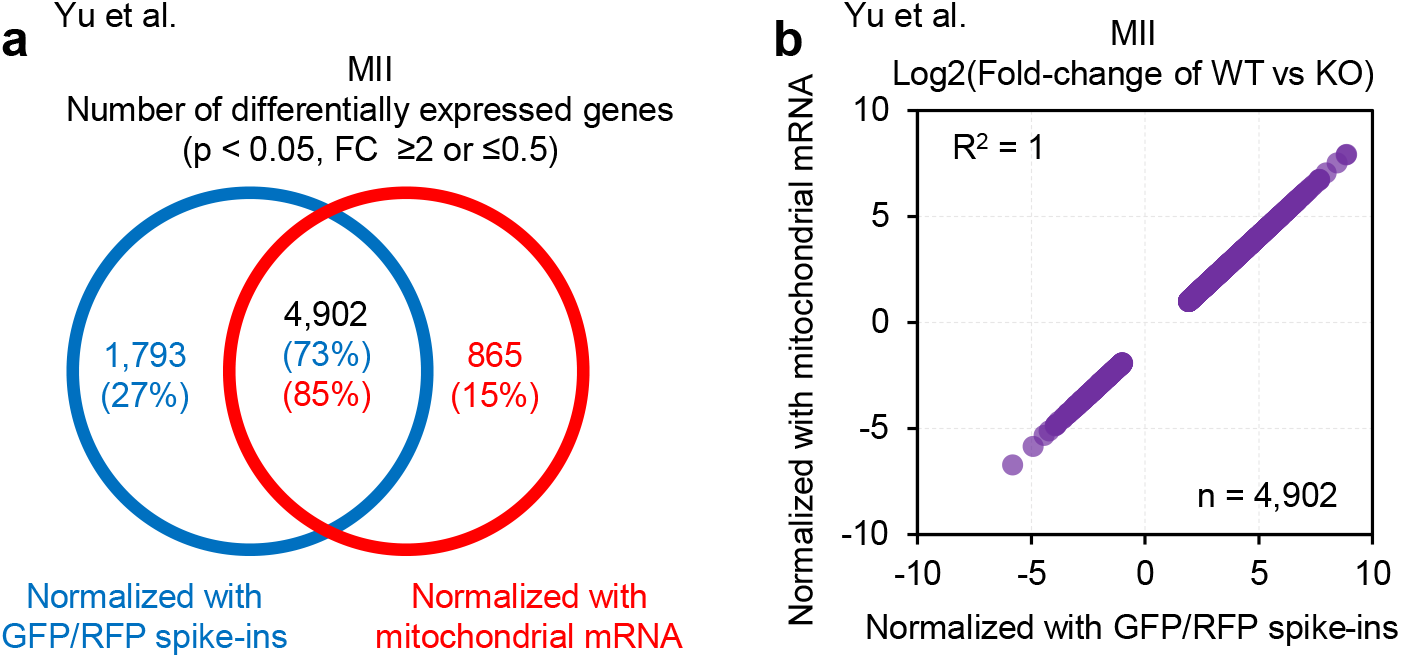
Comparison detection of differentially expressed genes using normalized read counts by exogenous spike-ins or mitochondrial mRNA. **a**, Venn Diagram showing differentially expressed genes (DEGs) in mouse *Btg4* KO MII oocytes^18^ quantified using read counts normalized by exogenous spike-ins (GFP and RFP mRNA) or mitochondrial mRNA. The number and percentage of DEGs are included in the diagram. **b**, Scatter plot of the expression changes of DEGs detected by counts from both normalization methods. Number of genes and Pearson’s correlation co-efficiency (R^2^) are included in the plot.

## Discussion

The normalization of RNA-seq data can remove systematic technical effects to ensure that technical bias has a minimal impact on analyzing expression levels. Sample variability is caused by several factors, including sample quality, the level of RNA yield, and user error. Typical normalization strategies mainly account for variability in sequencing depth and are based on the common assumption that the majority of the genes are not differentially expressed between samples. Control based normalization is developed for conditions that violate the above assumption to achieve better normalization, leading to the development of a set of external RNA, such as ERCC RNA spike-ins. However, adding exogenous spike-ins to RNA analysis experiments, especially for low input samples, is prone to technical variation in the percentage of reads mapping to the ERCC spike-ins between replicates^6^. In this study, we demonstrated that mitochondrial mRNA is a suitable endogenous reference for RNA-seq data from the same type of samples and the same library construction methods (Fig. 5, up).

**Fig. 5.**
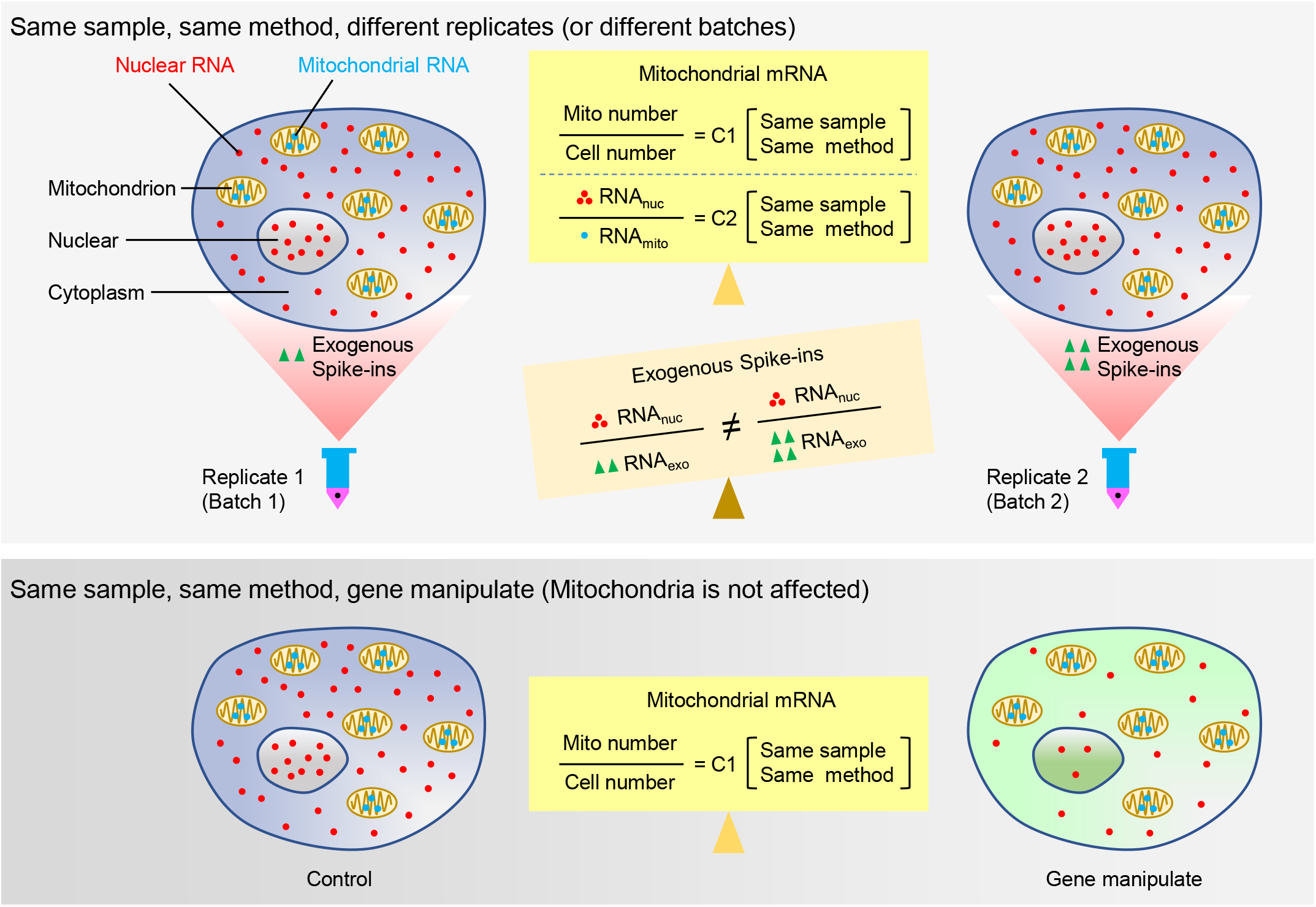
Summary of normalization of RNA-seq data using mitochondrial mRNA and exogenous spike-ins. For different replicates or different batches of the same sample, the ratio of the number of mitochondria to the number of cells is constant, which ensures a stable ratio of total nuclear mRNA to total mitochondrial encoded mRNA. This avoids problems caused by different amounts of exogenous mRNA spiked into the different replicates/batches. For treatments (including genetic or chemical treatments) that affect the global nuclear transcriptional level, the mitochondrial mRNA is enclosed in the mitochondria (which is typically not affected) and the ratio of mitochondria to the number of cells is constant. Therefore, normalization using mitochondrial encoded mRNA is a suitable method of correcting global bias at the transcriptome level.

Mitochondrial mRNA content for sample replicates is constant, meaning that mitochondrial mRNA can be used as a reference to judge the quality of RNA-seq data. In replicates from the same batch of samples following the same library construction procedures, significant differences in mitochondrial mRNA content suggest problems during the library preparation of this replicate (Extended Data Fig. 2).

Mitochondrial mRNA is also a suitable reference for comparing transcriptomes during gene manipulation or drug treatment, as long as it does not affect mitochondrial mRNA metabolism processes (Fig. 5, down). Mitochondria genomes (mitogenomes) are semi-autonomous organelles, meaning that transcription regulation is isolated from regulation in the nucleus. Therefore, most treatments that affect nuclear gene regulation do not affect mitochondria. For example, *Btg4* knockout, which prevents global deadenylation of nuclear encoded maternal mRNA, has no effect on mitochondrial mRNA deadenylation in either mouse oocytes or human zygotes (our mouse/human/mito paper). Therefore, a normalization strategy using mitochondrial mRNA as a reference can be used in most cases of biological treatment.

While using mitochondrial mRNA as a reference is convenient for RNA-seq data normalization, it cannot be used to compare samples of different cell types since the number of mitochondria in a cell can vary widely by organism, tissue, and cell type^17^. If the sample treatment affects mitochondrial gene expression, such as drug inhibiting mitogenome transcription, the mitochondrial mRNA cannot be used as a reference. In addition, if the purpose is to compare mitochondrial mRNA expression, using exogenous spike-ins is the option to choose.

While single-cell RNA-seq is a popular technique for cell typing and cellular heterogeneity analysis, bulk RNA-seq is the most commonly used technique for transcriptome analysis in most laboratories. Therefore, a simple and convenient method of transcriptome data normalization is powerful and of general interest. We believe that the simplicity and practicality of our novel normalization approach using mitochondrial mRNA as a reference will be widely adopted by many laboratories.

## Materials and Methods

### RNA-seq reads mapping

For the HeLa, HEK293T and mouse GV oocyte RNA-seq data, the raw sequencing data were downloaded from the SRA following their GEO accession. The raw reads were trimmed using trim_galore (https://github.com/FelixKrueger/TrimGalore). Then the data were mapped to the reference annotations (gencode.v38 from Ensembl for the human data, gencode.vM25 from Ensembl for the mouse data) and quantified using salmon^22^. For the zebrafish data, the read counts for each gene of the zebrafish reference genome (Zv9, Ensembl, v.67) and the ERCC spike-ins were downloaded from the Bioconductor (zebrafishRNASeq, 10.18129/B9.bioc.zebrafishRNASeq)^5^. Read counts of the 13 mitochondrial protein-coding genes (*Nd1, Nd2, Nd3, Nd4, Nd4l, Nd5, Nd6, Cox1, Cox2, Cox3, Atp6, Atp8*, and *Cytb*) were treated as mitochondrial mRNA.

### Exogenous spike-ins normalization

For ERCC spike-ins normalization, the ERCC sequences (ERCC92.fa) were downloaded from Thermofisher (https://www.thermofisher.cn) containing the 92 polyadenylated RNAs in ERCC spike-ins. The ERCC reads were counted by mapping the samples to the ERCC sequence by bowtie2^23^. For the zebrafish data, the ERCC counts were included in the zebrafishRNASeq dataset (10.18129/B9.bioc.zebrafishRNASeq)^5^. The spike_in normalized read counts for each gene was calculated as 1,000,000*gene_fragment_counts/ERCC_read_counts.

### Mitochondrial mRNA normalization

The mitochondria mRNA normalized read counts for each gene was calculated as 1,000,000*gene_fragment_counts/mito_mRNA_read_counts.

## Data availability

This study includes analysis of the following published data: RNA-seq data of Di et al. (Genome Sequence Archive database (GSA) accession no. CRA002081), Rong et al. (Gene Expression Omnibus database (GEO) accession no. GSE135787), Zhao et al. (GEO accession no. GSE139072), Yu et al. (GEO accession no. GSE71257), and Risso et al. (GEO accession no. GSE53334).

## Acknowledgements

This work was supported by the National Key Research and Development Program of China (2018YFA0107001), the Strategic Priority Research Program of the Chinese Academy of Sciences (XDA24020203), National Natural Science Foundation of China (31970588), Natural Science Foundation of Heilongjiang province (YQ2020C003), the China Postdoctoral Science Foundation (2020M670516, 2020T130687), and the State Key Laboratory of Molecular Developmental Biology.

## Author Contributions

Yusheng Liu, Falong Lu and Jiaqiang Wang conceived the project and designed the study. Yusheng Liu, Yiwei Zhang, Falong Lu and Jiaqiang Wang analyzed the sequencing data. Yusheng Liu and Jiaqiang Wang organized all figures. Yusheng Liu, Falong Lu and Jiaqiang Wang supervised the project. Yusheng Liu, Falong Lu and Jiaqiang Wang wrote the manuscript with the input from the other authors.

## Figure legends

**Extended Data Fig. 1.**
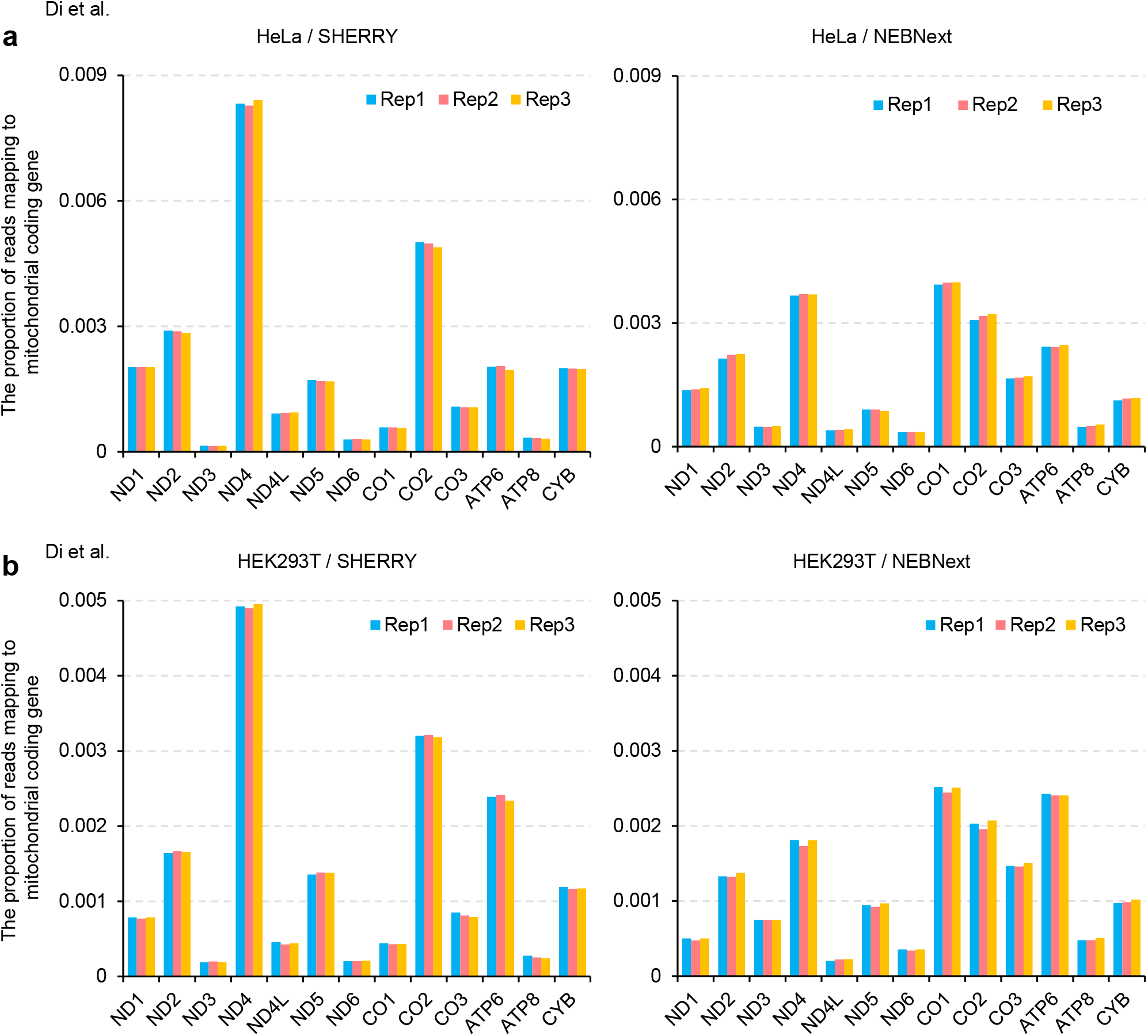
Level of individual mitochondrial coding gene in multiple RNA-seq datasets. Proportion of read counts mapped to each mitochondrial coding gene in HeLa cells (**a**) and HEK293T cells (**b**). Three replicates from the datasets are shown as bar plots. Two different RNA-seq library constructing methods, SHERRY (left) and (right) NEBNext, are used in these datasets^20^.

**Extended Data Fig. 2.**
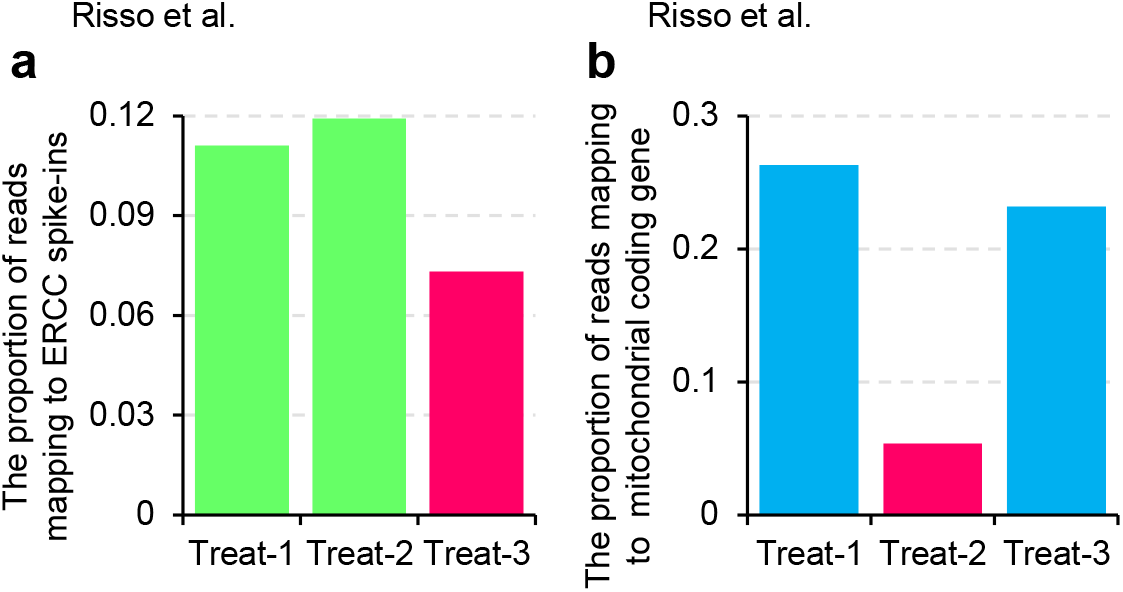
Level of exogenous spike-ins or mitochondrial mRNA in a group of zebrafish treatment dataset of three replicates. Bar plots indicate the proportion of total reads mapping to the ERCC spike-ins (**a**) or mitochondrial mRNA (**b**) in a zebrafish treatment dataset of three replicates^5^. The bars in red indicate samples with significant deviations from the other two replicates.

